# Downstream effects of dre-miR-210-5p in zebrafish primary ovarian cell culture

**DOI:** 10.1101/2023.08.10.552851

**Authors:** T.A. van Gelderen, L. Ribas

**Affiliations:** Institut de Ciències del Mar, Consejo Superior de Investigaciones Científicas (ICM-CSIC), 08003, Barcelona

## Abstract

MicroRNAs (miRNAs) are small, non-coding RNAs that are involved in post-transcriptional gene regulation in many cellular functions and are highly conserved throughout evolution. In teleost fish species, miRNAs are believed to play a role in the reproductive system, but more research is needed to better understand the functions of miRNAs in fish gonads. Furthermore, miR-210 has previously been described to be involved in many processes, such as hypoxia response, angiogenesis, cell proliferation and male infertility. The aim of this study was, first, to develop an *in vitro* model in fish to study the functions of miRNAs and second, to identify target genes of dre-miR-210-5p by establishing a primary ovarian cell culture in zebrafish (*Danio rerio*). The cell culture was performed by isolating ovaries from adult female fish (n=4) which were incubated with dre-miR-210-5p mimic or scramble miRNA mimic. Cell survival was studied by flow cytometry analysis and fluorescent microscopy. Following, the effect of dre-miR-210-5p on ovarian cells was uncovered by RNA-sequencing, identifying ∼6,000 targeted-genes differentially expressed (DEGs), of which ∼2,600 downregulated and ∼3,400 upregulated. GO term and KEGG pathway analyses showed downregulated genes involved in cell cycle processes and reproduction-related pathways while, in contrast, immune-related pathways were upregulated after miR-210 mimic treatment in the ovarian cells. To provide the molecular mechanisms triggered by miRNA-210, seed regions of targeted genes were identified and *in silico* analysis classified DEGs as potential biomarkers in reproduction or immune cell types. These results support the crosstalk between the reproductive and immune system in which dre-miR-210-5p plays a key role in the transcriptomic alteration in the fish ovaries.

**Take home message:** miR-210 promotes the immune response and suppresses oocyte meiosis in the ovarian cells in zebrafish

## Introduction

Micro RNAs (miRNAs) are small, non-coding RNAs involved in post-transcriptional gene regulation. The post-transcriptional regulation is one type of the three main epigenetic mechanisms and so, the role of the miRNA in the cellular response is relevant to better understand the epigenetic mechanisms. In humans, miRNAs are seen as an opportunity to identify novel candidates for genetic treatments or for marker diseases (Condrat *et al*., 2020). In fish, in the last years, the interest of miRNAs is growing exponentially, and for example, public and specific miRNA database for fish have been developed (Kozomara, Birgaoanu and Griffiths-Jones, 2019; Desvignes *et al*., 2022)

miRNAs are involved in many cellular processes. They are critical regulators of fish development and morphogenesis by controlling the expression of genes involved in embryonic development, tissue differentiation, and organ formation (Herkenhoff *et al*., 2018). miRNAs are also involved in regulating immune responses in fish by modulating the expression of immune-related genes, including those involved in pathogen recognition, immune cell activation, cytokine production, and inflammation (Andreassen and Høyheim, 2017). Furthermore, miRNAs are involved in the reproductive system, for example, during sex differentiation in the tiger pufferfish (*Takifugu rubripes*) (Yan *et al*., 2021). In adulthood, specific miRNAs were found in the ovaries and testes (Bhat *et al*., 2021). For example, miR-202-5p was essential for oocyte proliferation and female fertility in Japanese medaka (*Oryzias latipes*) (Gay *et al*., 2018), and their expression changed significantly throughout follicle development (Wong *et al*., 2018). Environmental influences, such as temperature, salinity, oxygen and feed caused alterations in the miRNA expression patterns (Cao *et al*., 2023). Recently, miRNAs have been identified as heat-recorders by long-term changes in zebrafish (*Danio rerio*) heat-treated gonads (van Gelderen *et al*., 2022), as well as in Atlantic cod (*Gadus morhua*) (Bizuayehu *et al*., 2015) and European sea bass larvae (Papadaki *et al*., 2022).

miR-210 responds to hypoxic environmental stress by upregulating its expression when fish are exposed to low oxygen levels. The regulation is activated by the binding of hypoxia inducible factor 1 alpha (*HIF1*α) to the hypoxia responsive element (HRE) on the miR-210 promoter (Huang *et al*., 2009). During hypoxia, miR-210 is involved in many biological processes, such as angiogenesis, DNA replication and cell division (Ivan and Huang, 2014), but also in normoxia conditions (Bavelloni *et al*., 2017). Its role in the sexual organs has been associated with male fertility (Duan, Huang and Sun, 2016; Tang *et al*., 2016) and with correct testicular development (Elias *et al*., 2022). In the female sex organ, however, its function remains relatively unknown. Moreover, miR-210 also participates in tissue regeneration and development in fish. It has been shown to promote angiogenesis, which is the formation of new blood vessels, by targeting specific genes involved in vascular growth (Yang *et al*., 2016). This function is crucial for fish during tissue repair and regeneration processes, ensuring efficient nutrient and oxygen supply to the renewing tissues (Cooke and Kalluri, 2008). Furthermore, miR-210 regulated the expression of genes involved in the inflammatory response in fish. It can modulate the production of pro-inflammatory cytokines, such as interleukin-1β (IL-1β) and tumor necrosis factor-alpha (TNF-α), which are crucial for initiating and regulating the inflammatory cascade during immune responses (Su *et al*., 2021). By controlling the expression of these genes, miR-210 can influence the intensity and duration of the inflammatory response and regulate the differentiation and activation of immune cells, such as macrophages (Chan *et al*., 2009). Thanks to the pleiotropic role of miR-210 in the immune and the reproduction system, it makes a suitable candidate to deeply study the interactions between these two systems in fish.

Understanding the functions of miRNA and identifying their direct targets has presented a considerable challenge (Krützfeldt, Poy and Stoffel, 2006). Currently, different methods exist to identify target genes and miRNA function. With luciferase assays, the binding capability of a miRNA to the 3’UTR of a gene can be determined in a simulated environment, using a plasmid containing the 3’UTR sequence of the gene of interest alongside a fluorescent protein (Tomasello, Cluts and Croce, 2019). Two techniques offer the possibility to precipitate the bound miRNA-mRNA complex, namely pull-down assay and CLIP (Cross-Linking and Immunoprecipitation), by either introducing biotinylated RNA molecules, designed to mimic the miRNA, that bind to the mRNA (Dash *et al*., 2018), or cross-linking and immunoprecipitation of the miRNA-mRNA complex (Wen *et al*., 2011). These methods, however, do not inform on the general function of a miRNA in a cellular environment. Knockout by CRISPR (Chang *et al*., 2016) or knockdown by miRNA inhibitor transfection (Robertson *et al*., 2010) of a miRNA can show critical functions, similarly to overexpression by miRNA mimic transfection (Wang, 2011). Even though CRISPR offers an *in vivo* solution, crossings may take many trials, and depending on the species, maturing of the animal may be a lengthy process. In the last years, a novel method by miRNA inhibitors and mimics *in vitro* cell cultures offers an immediate time- and dose-dependent solution. It is based on introducing a synthetic endogenous (mimic) or complementary (inhibitor) miRNA sequence to the cell, causing either higher miRNA activity or suppressing miRNA counts.

When searching the number of publications in the Web of Science and typing in the topic search “miRNA mimic human” yields ∼1.850 results; “miRNA mimic mouse” yields ∼1.150 results and most notably; “miRNA mimic fish” only yields 45 results. Thus, the amount of available data in fish studying miRNA function by mimic strategies are scarce. For example, mimic studies have been performed in macrophage cell cultures in miiuy croaker (*Miichthys miiuy*) (Chu *et al*., 2017, 2019; Xu *et al*., 2018) and in ZF4 cells in zebrafish (Ji *et al*., 2020). However, to our knowledge, no studies regarding fish gonadal cells have ever been performed. In order to obtain insight into the function of dre-miR-210-5p in an ovarian cellular environment, we present a protocol for miRNA mimic incubation in primary ovarian cell culture in zebrafish. To validate our method, cell survival was studied by flow cytometry and fluorescent microscopical analysis, and assessment of the post-transcriptional regulation of the mimic was performed by RNA-sequencing. Furthermore, target DEGs and seed regions were identified in those altered pathways and potential biomarkers for the cultured cell types were identified. In this study, downstream effects of miR-210-5p in zebrafish ovarian cells were described for the first time.

## Materials and methods

### Fish housing

Zebrafish (TU strain) were housed in a commercial rack (Aquaneering, San Diego, CA, USA) in an *ad hoc* chamber facility with a recirculating water system. The chamber had a regulated photoperiod (12h dark; 12h light), an air temperature of 26 ± 1°C and a humidity of 50 ± 3%. The water quality parameters were monitored every day and were as follows: a conductivity between 750 and 900 µS, dissolved oxygen between 6.5 and 7 mgL^-1^, a pH of 7.2 ± 0.2 and a water temperature of 28 ± 0.2°C. Ammonia, nitrate, and nitrite water levels were checked weekly by a commercial colorimetric test (Test kit Individual Aquarium Water Test Kit, API, United Kingdom). The fish were fed three times a day with commercial powdered food (AquaSchwarz, Göttingen, Germany). This food was supplemented with *Artemia nauplii* (AF48, INVE Aquaculture, Dendermonde, Belgium). The tanks were cleaned three times a week to avoid debris at the bottom and walls of the tanks.

### Ovarian cell culture

Four adult female zebrafish were sacrificed using cold thermal shock and their ovaries were removed immediately and transferred to a plate (Figure 1). The ovaries were disinfected with 70% ethanol for 5 seconds and washed with phosphate buffered saline (PBS). Based on the ovary weight, samples were incubated with 500, 600 or 700 µL of 1 mg/mL collagenase in L-15 medium on a shaker for 1 hour at room temperature (RT). In order to stop the reaction and to create a homogenous cell suspension, the cell/collagenase mix was diluted 1:10 in L-15 medium supplemented with 10% Fetal Bovine Serum (FBS) and 1% penicillin/streptomycin. For each fish sample, the cell suspension was pipetted in quadruplicates for each of the four experimental groups: miR-210 mimic (Group 1), scramble mimic (negative control, Group 2), Transfection Reagent (TR, Group 3) and cell suspension (CS, Group 4) on a 24-wells plate with a volume of 500 µL per well. For fluorescence microscopy and flow cytometry analyses, 300 µL was used to assess cell survival (T0).

**Figure 1.**
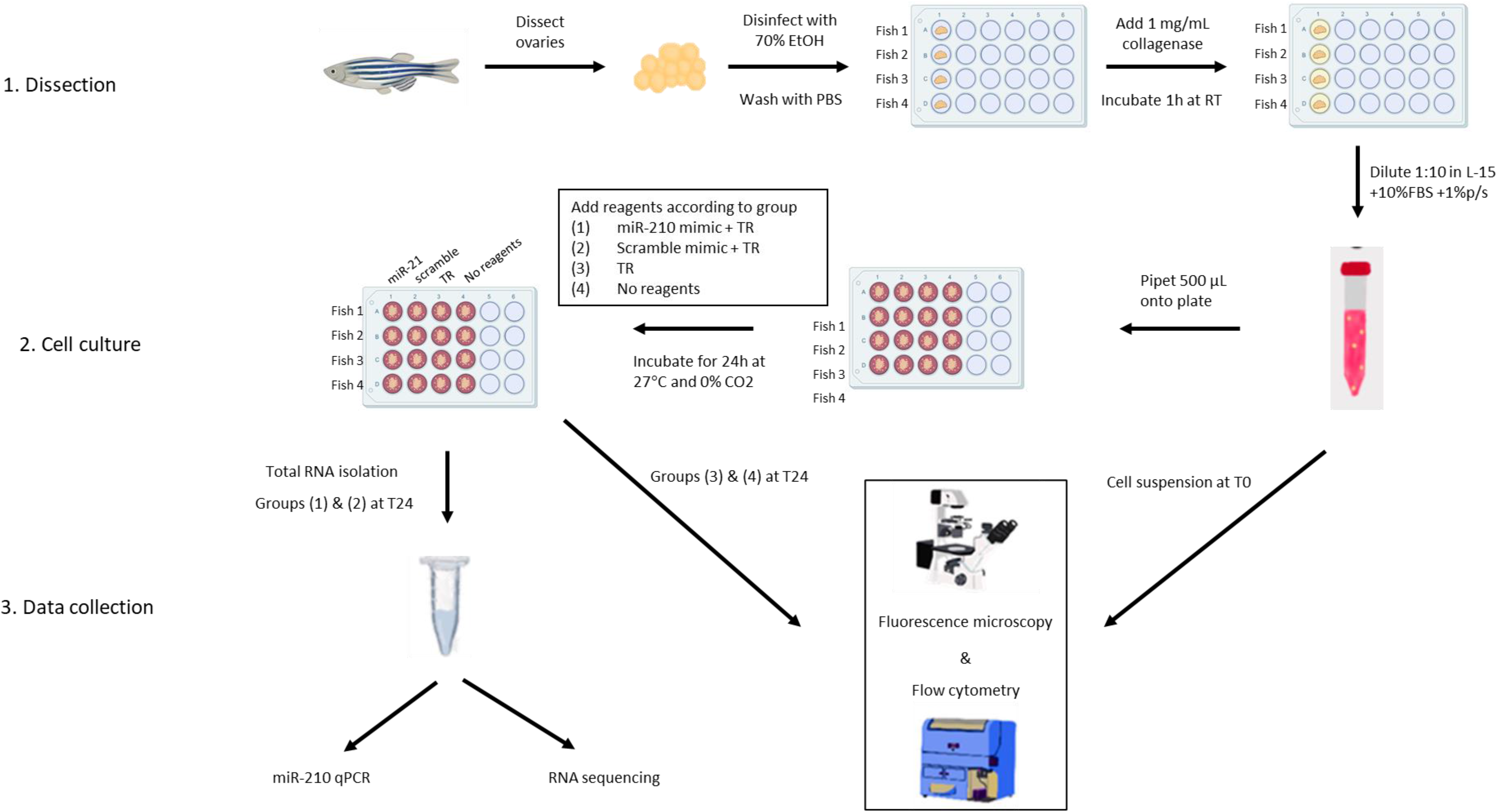
Schematical overview of the materials and methods of miRNA mimic incubation on a primary ovarian cell line.

### Flow cytometry

In order to assess the cell survival after 24h (T24), for each fish, 300 µL ovarian cells at T0 and the TR-group 3 and CS-group 4 at T24 were transferred to 1 mL flow cytometry vials. Then, the samples were incubated for 10 min with 1% SybrGreen and 1% propidium iodine (PI). In a Cytoflex S flow cytometer (Beckman Coulter), cells were passed for 120 seconds at 30 µL per minute. Cell damage was assessed by PI intensity and DNA content by SybrGreen. Cell populations were determined based on cell population patterns and cell counts were compared between T24 and T0 to determine cell survival.

### Fluorescence microscopy

Ovarian cells were fixed over-night (O/N) with 20% glutaraldehyde at 4°C in a 1.5 mL Eppendorf tube. Cellulose acetate filter of 0.8 μm (AA, 25 mm) were placed in a tower filtration system with a 0.2 μm polycarbonate black filter (25 mm) on top. The samples were added to individual filters alongside 1 µL of 0.5 mg/µL DAPI and 1 µL of 1:100 Wheat Germ Agglutinin (WGA) and incubated for 10 minutes. Then, using the filtration system, the filters were dried. The filters were then placed on a slide and covered with immersion oil and a cover slip.

### Mimic incubation

In unsupplemented L-15 medium, 3 µg (10.7 µL) of double stranded miRNA mimic (5’ AGCCACUGACUAACGCACAUUG 3’) or negative control (5’ UUCUCCGAACGUGUCACGUTT 3’) purchased from GenePharma (Shanghai, China) was diluted together with 6 µL of Roche X-tremeGene siRNA Transfection Reagent (TR) to a final volume of 50 µL, after which they were incubated for 15 min at RT. Then, the miR-210 mimic/TR mixture, scramble mimic/TR mixture or unsupplemented L-15 medium was added to the zebrafish ovary cell suspension previously prepared for each group. Additionally, 6 µL of TR without miRNA mimic was added to the ovarian cell culture of group (3). Following, ovarian cells were incubated for 24h at 27°C.

### RNA extraction

After 24h, cell suspensions were transferred to 15 mL falcon tubes and spun down at 3000g for 10 minutes. In order to wash the cells adhered to the plate, 200 µL PBS was added. The supernatant of the suspended cells was removed and PBS from the plate plus 300 µL clean PBS was added to each corresponding sample. Cells were spun down again and supernatant was removed. To each well, 150 µL RLT buffer (Qiagen) with 1:10 β-mercaptoethanol (55mM) was added and incubated for 10 minutes. A total of 200 µL plus 150 µL from each corresponding well was added to the cell pellets, after which they were vortexed for 1 min. RNA extraction was followed according manufacturer’s instructions. RIN values were assessed using BiOptic Qsep100 and RNA yield, 230/260 and 260/280 values were determined using Nanodrop (Thermo Scientific). Samples with a RIN >7.5, concentration >100 ng/µL, 260/280 >1.8 and 230/260 >1.8 were selected for RNA sequencing.

### miR-210 transfection validation

To check successful transfection of miR-210 mimic in the ovarian cell samples, we performed miRNA cDNA synthesis and measured miR-210 overexpression by qPCR. cDNA was synthesized from four biological replicates incubated with either miR-210 mimic or scramble mimic, by following the manufacturer’s instructions of the high specificity miRNA 1st-strand cDNA synthesis kit (Agilent Technologies). First, the polyadenylation step was performed with 100 ng RNA for each sample. The mixture was incubated at 37°C for 30 minutes, then at 95°C for 5 minutes and immediately transferred to ice. Of the polyadenylation mixture, 4 µL was used for the 1st-strand cDNA synthesis. The synthesis reaction was incubated at 55°C for 5 min, 25°C for 15 min, 42°C for 30 min and finally, to terminate reverse transcription, 95°C for 5 min. This product was directly used for qPCR, using the dre-miR-210-5p as the forward primer: 5’ AGCCACTGACTAACGCACATTG 3’ and the Universal Reverse Primer (N° 600037, Agilent Technologies) as reverse primer.

### RNA sequencing

For each fish, a library was constructed from primary zebrafish ovarian cell culture for the two studied groups (N=8 total, N=4 for each group). DNBSEQ Eukaryotic Strand-specific mRNA libraries were prepared at external services (BGI genomics, Hong Kong). Sequencing (DNBSeq) was performed at paired-end mode with a read length of 100 bp by BGI Genomics in Hong Kong. Internal filtering was done by BGI Genomics using SOAPNuke software (Chen *et al*., 2018). Data filtering includes removing adaptor sequences, contamination and low-quality reads from raw reads. After filtering, sequencing yielded an average of 34 million reads per sample. All samples had a QC20 of > 98% and a GC count around 49%.

### Bioinformatics

Sequenced libraries were aligned by Hisat2 (Kim *et al*., 2019) to the zebrafish genome (GRCz11) and reads were counted by FeatureCounts (Liao, Smyth and Shi, 2014). Differential expression of the genes was determined using EdgeR (Robinson, McCarthy and Smyth, 2009), upholding a threshold of FDR<0.05. To visualize similarity in expression patterns between the samples, a Multi-Dimensional Scaling (MDS) plot was created with EdgeR (Robinson, McCarthy and Smyth, 2009). Based on the origin of the samples, for the MDS analysis, paired samples and batch effect was considered. A heatmap of the differentially expressed genes (DEGs) were made using the pheatmap (version 1.0.12) R package. Enriched GO terms and KEGG pathways were calculated using ClusterProfiler (Yu *et al*., 2012) and KEGG graphs created with pathview (Luo and Brouwer, 2013).

### Violin plot

A violin plot was made using the ggplot2 package in R (Wickham, 2016). Cell marker genes were extracted from Liu *et al*. (Liu *et al*., 2022) and Jiang *et al*. (Jiang *et al*., 2021), joined and assigned to one of the following categories: innate immune cell, granulocyte, lymphoid cell, myeloid cell, pre-oocyte, oocyte, somatic cell, vasculature cell and epithelial cell. Then, the DEGs identified in our study were classified by one of the cell categories. When no category was able to be assigned, it was classified as “other”.

### RNA-seq validation

Validation of the RNA sequencing data was done by qPCR for 8 DEGs for each fish individually. In order to increase biological significance, besides two samples used in RNA sequencing, two additional experimental samples were used. cDNA was generated using the Roche Transcriptor First Strand cDNA Synthesis Kit (Cat# 04379012001) with 100 ng RNA as input, following manufacturer’s instructions. qPCR was performed using the qPCRBio SyGreen blue mix low ROX (PCR Biosystems). A mix of 5 µL 2x qPCRBIO SyGreen Blue mix, 0.4 µL forward primer, 0.4 µL reverse primer, 1 µL cDNA and H2O up to 10 µL was made for each sample. Samples were incubated at 95°C for 2 minutes, followed by 40 cycles of 95°C for 30 seconds and 60°C for 30 seconds, after which the melting curve was determined by incubating at incrementing temperatures from 60 to 65°C by 0.5°C steps and finally denatured at 95°C for 1 minute. Primer sequences for the selected miRNA-target genes are shown in Suppl Table I.

## Results

### Ovarian cell survival

Cell populations of the primary ovarian cell cultures were assessed by flow cytometry and fluorescence microscopy at T0 and T24. We identified three main populations, pink (main cell population), purple (main subpopulation) and green (damaged cells). Figure 2 shows an example of the cell patterns found in the flow cytometry analysis at the two sampling times and with or without TR. The three patterns for the PI intensity and DNA content were mostly the same, indicating that cell survival was relatively high after the incubations.

**Figure 2.**
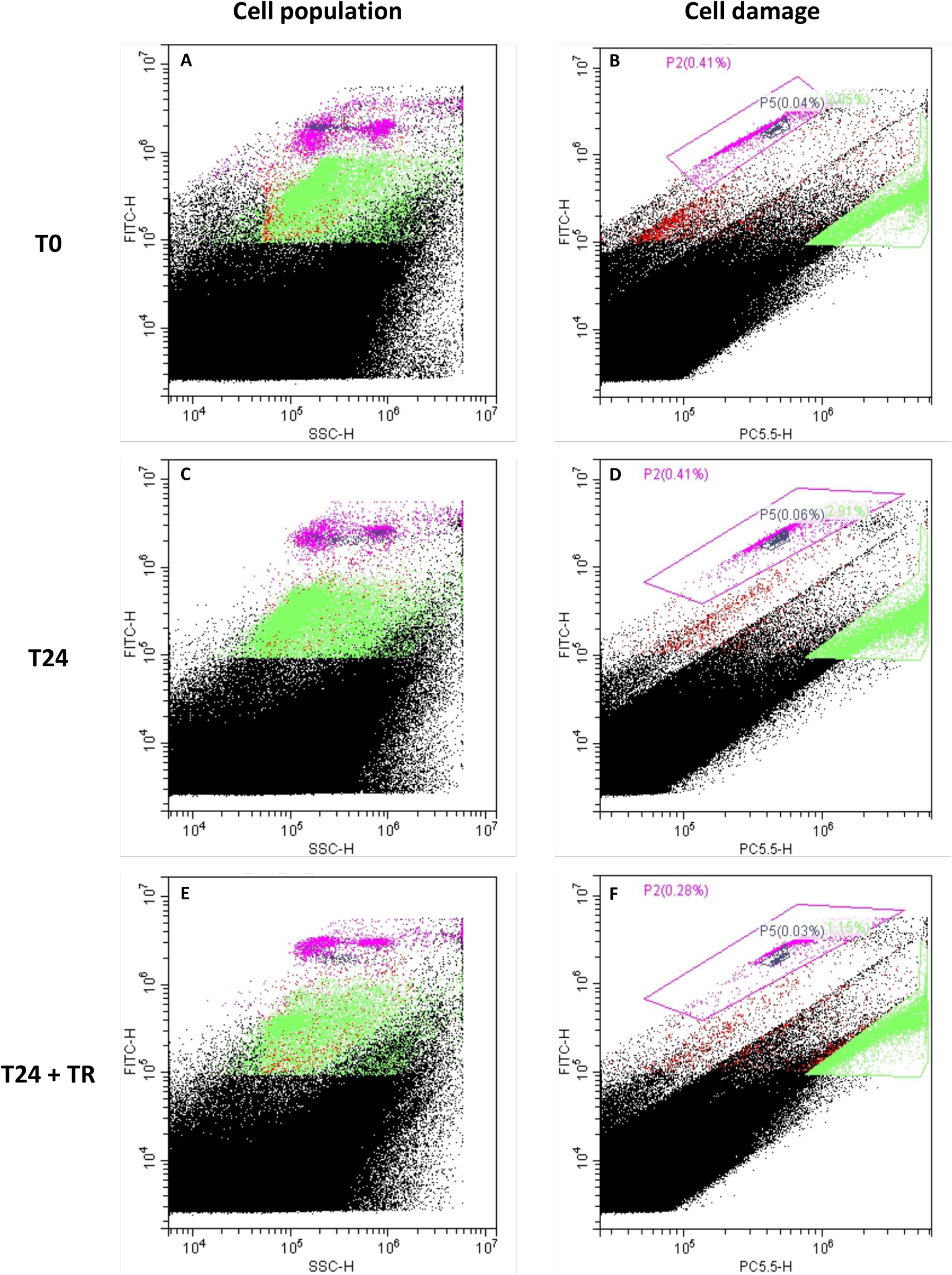
Flow cytometry of zebrafish ovarian cell culture at T0 (A, B), T24 (C, D) and T24 after incubation with transfection reagent (E, F). (A, C, E) show side scatter against DNA amount, measured by DAPI, whereas (B, D, F) show cell damage by incubation with propidium iodide against DNA amount. Populations marked in purple are believed to be main ovarian cells, population marked in green are damaged cells.

Furthermore, after measuring the cell concentration by flow cytometry at T0 of cell population P2 (pink), concentrations were adjusted to ∼12,000 cells/mL on the 24-wells plate. After 24 hours, cell concentration was measured of the cell suspension (group 4) and CS + transfection reagent (group 3) in order to imitate miRNA mimic conditions. Samples containing TR had lower concentrations compared to CS, namely 82%, 90%, 50% and 29% for each of the four fish used, respectively (Table I), given an average of 62% of cell survival T24 after the treatment.

**Table 1.** Cell counts of primary ovarian cell culture in flow cytometry at time 0 and 24 hours of incubation with or without transfection reagent (TR). Percent of the cell counts at T24 between groups with or without TR are given

| Fish number | T0 | T24 | T24+TR | %TR |
| --- | --- | --- | --- | --- |
| 1 | 175.892 | 117.867 | 97.011 | 82 |
| 2 | 145.390 | 118.978 | 107.161 | 90 |
| 3 | 89.082 | 145.945 | 73.175 | 50 |
| 4 | 68.226 | 115.191 | 33.330 | 29 |

The shape and size of the cultured cells were evaluated by fluorescence microscopy after 24h with TR incubation. Different stages of oocyte maturation were observed (Figure 3A-D) with no visual significant differences between the T0 and the two groups at T24 (group 3-TR and group 4-CS). Four oocyte stages were observed (Selman *et al*., 1993): Stage I: primary growth stage (follicle diameter 7-140 µm, Figure 3A), Stage II: cortical alveolus stage (follicle diameter = 140-340 µm, Figure 3B), Stage III: vitellogenesis (follicle diameter = 340-690 µm, Figure 3C) and Stage IV: oocyte maturation (follicle diameter = 690-730 µm, Figure 3D).

**Figure 3.**
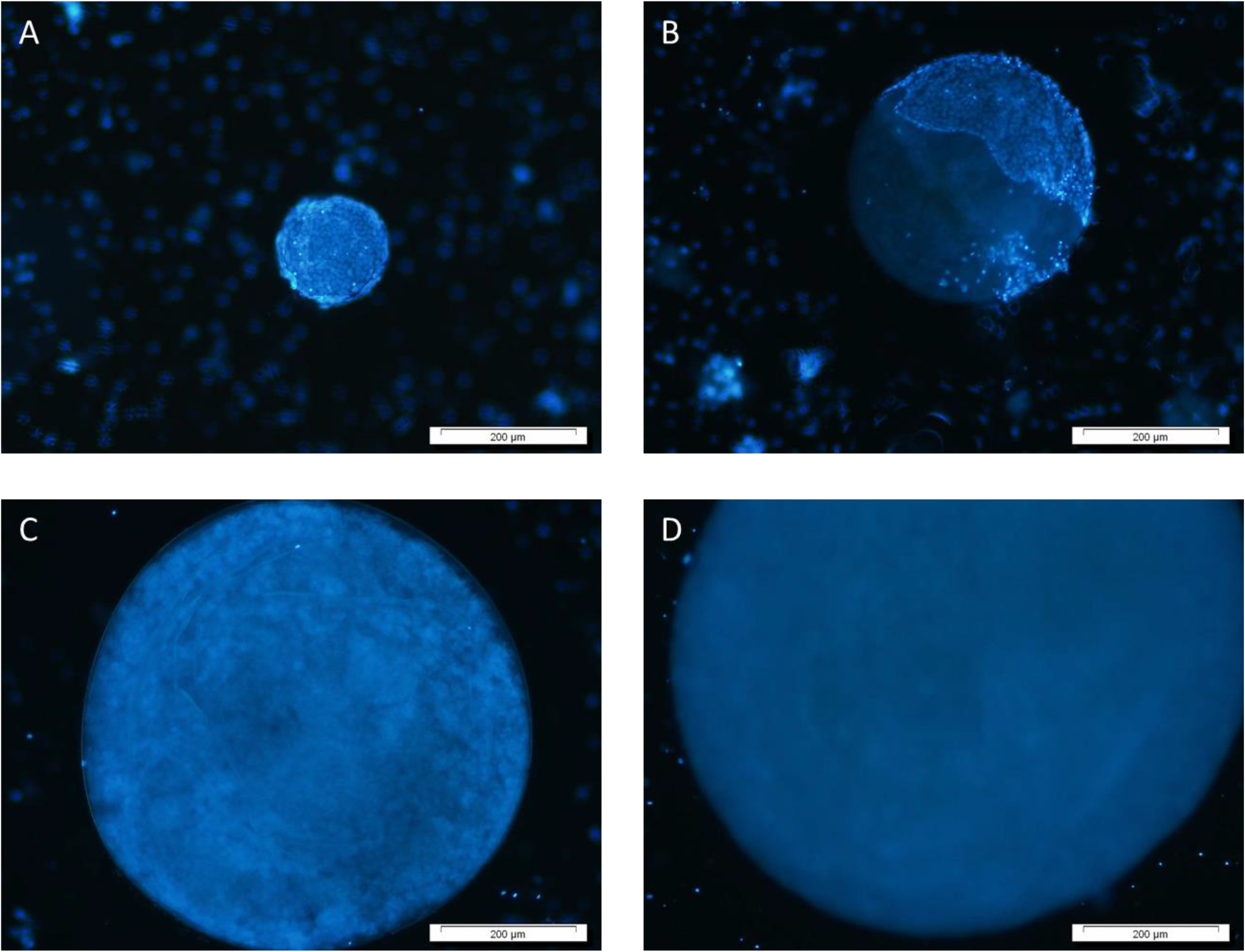
Fluorescence microscopy of oocytes of different maturation stages in a primary ovarian zebrafish cell culture. A. Stage I: primary growth stage (follicle diameter 7-140 µm) B. Stage II: cortical alveolus stage (follicle diameter = 140-340 µm. C. Stage III: vitellogenesis (follicle diameter = 340-690 µm). D. Stage IV: oocyte maturation (follicle diameter = 690-730 µm).

### miRNA mimic incubation

After 24h incubation of primary ovarian cell culture with dre-miR-210-5p and scramble mimics, a 3.4-fold upregulation was measured in the miRNA of interest mimic samples (Suppl Figure S1A). This data showed the presence of the miR-210 in ovarian cell cultures after the treatments.

### RNA sequencing

RNA sequencing yielded an average of 34 million sequences per library being a total of 684 million sequences. After trimming and alignment to the zebrafish genome, 32,520 transcripts were identified. After filtering, 21,545 transcripts remained for differential expression analysis performed with EdgeR (Robinson, McCarthy and Smyth, 2009). An MDS analysis, considering the paired samples and removing batch effect, was performed. Results clustered the samples based on treatment or control, with a dimension of 68% on the x-axis and 18% on the y-axis (Suppl Figure S1B). After differential expression analysis, 5,894 genes were identified with a false discovery rate (FDR) <0.05, of which 3,373 and 2,521 were up- or downregulated, respectively (Dataset 1). Notably, 16 genes, or close orthologues, that were significantly differentially regulated in our data have previously been described as direct targets of miR-210 (Table II). The relative expression pattern of each sample of the total DEGs was determined and visualized by a heatmap, showing high consistency of the differential expression patterns between biological replicates (Figure 3). Raw sequencing data was submitted to the Sequence Read Archive (SRA) from NCBI (https://www.ncbi.nlm.nih.gov) with the accession number: SUB13752204 (made public upon publication).

**Table 2.**
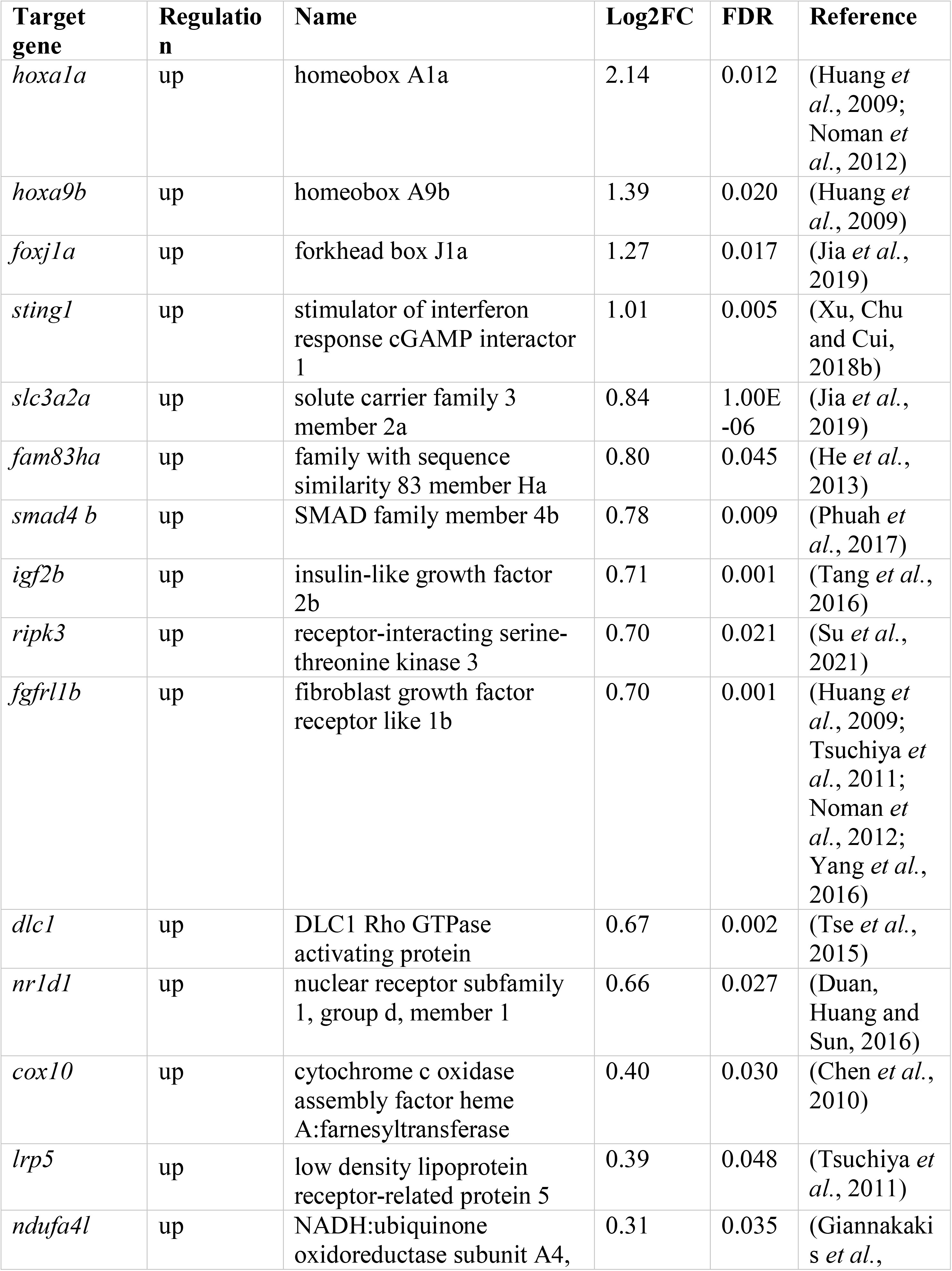
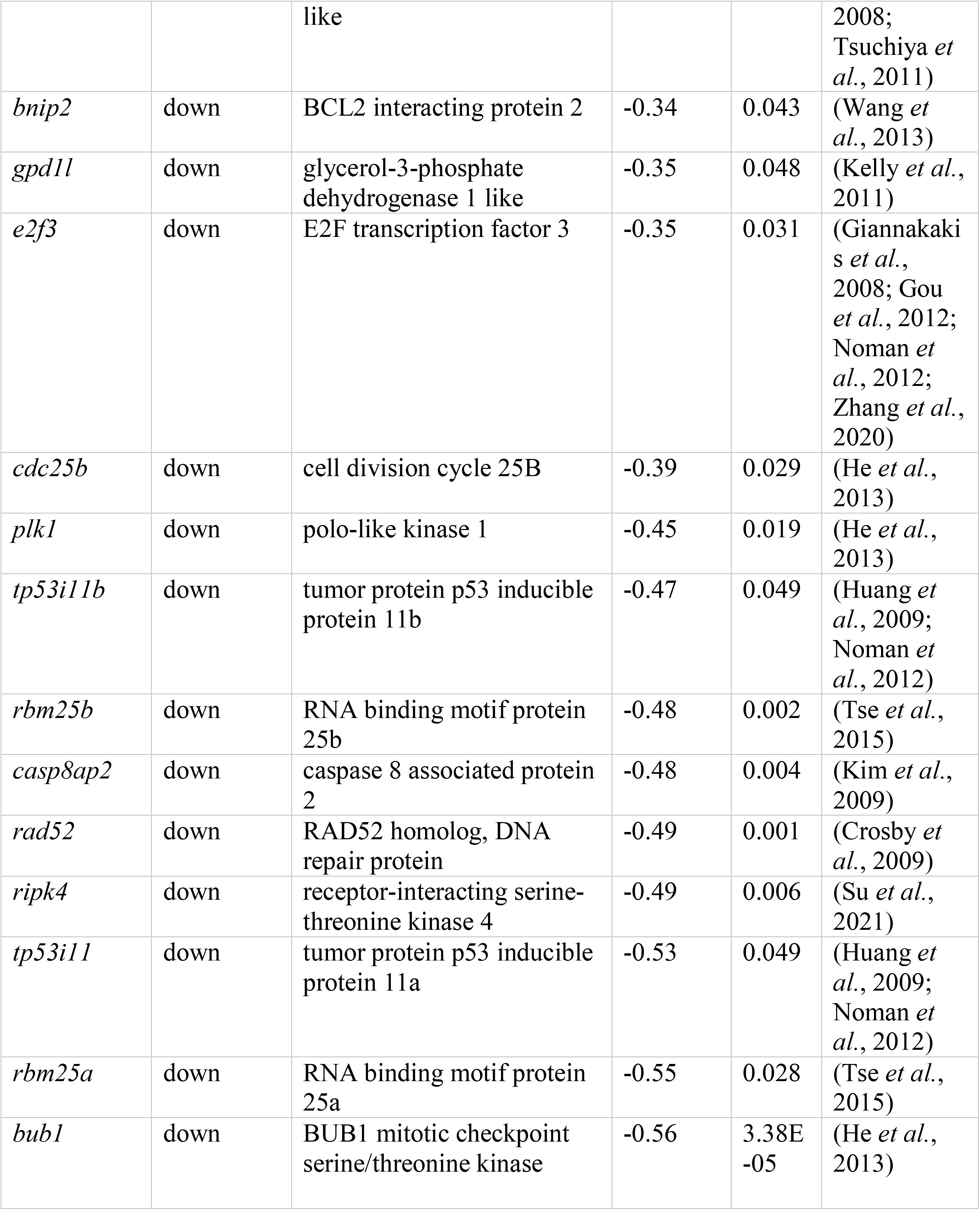
List of target genes of dre-miR-210-5p identified in a primary ovarian cell culture in zebrafish (*Danio rerio*)

### RNA-seq validation

RNA sequencing data was validated by qPCR by measuring eight DEGs, resulting in an R^2^ of 0.8759 and a *P-value* of 0.00063 (Suppl Figure S3). For the validation data, a total of four animals per group were used, being two of the used samples from fish not included in RNA sequencing data, thus, the validation of the technique was conclusive.

### GO and KEGG

In order to discover the function of the up- and downregulated genes, enriched GO terms and KEGG pathways were explored. Genes belonging to immune-related cells were shown to be mainly upregulated, as well as blood vessel-related genes and ovarian supporting cells. On the other hand, germ cell-related genes were mainly downregulated.

For the upregulated genes, 383 enriched GO terms were identified (adjusted *p*-value<0.05) (Dataset 2). Notably, most GO terms were related to the immune system and angiogenesis. The top 15 most significant (adjusted *p*-values 2.47E-28 to 1.58E-09) GO terms were shown in Figure 4A of which 13 were immune-related. For example, “immune response” (170 genes), interspecies interaction (137 genes) and “response to biotic stimulus” (130 genes). Furthermore, 32 KEGG pathways were identified as significantly enriched (adjusted *p*-value<0.1) (Figure 4B). The most significantly enriched immune-related pathway, namely “Phagosome”, was highlighted in Supplementary Figure S3 to dissect the alteration of this pathway after miR-210 treatment. This pathway contains 66 genes differentially expressed in our data.

**Figure 4.**
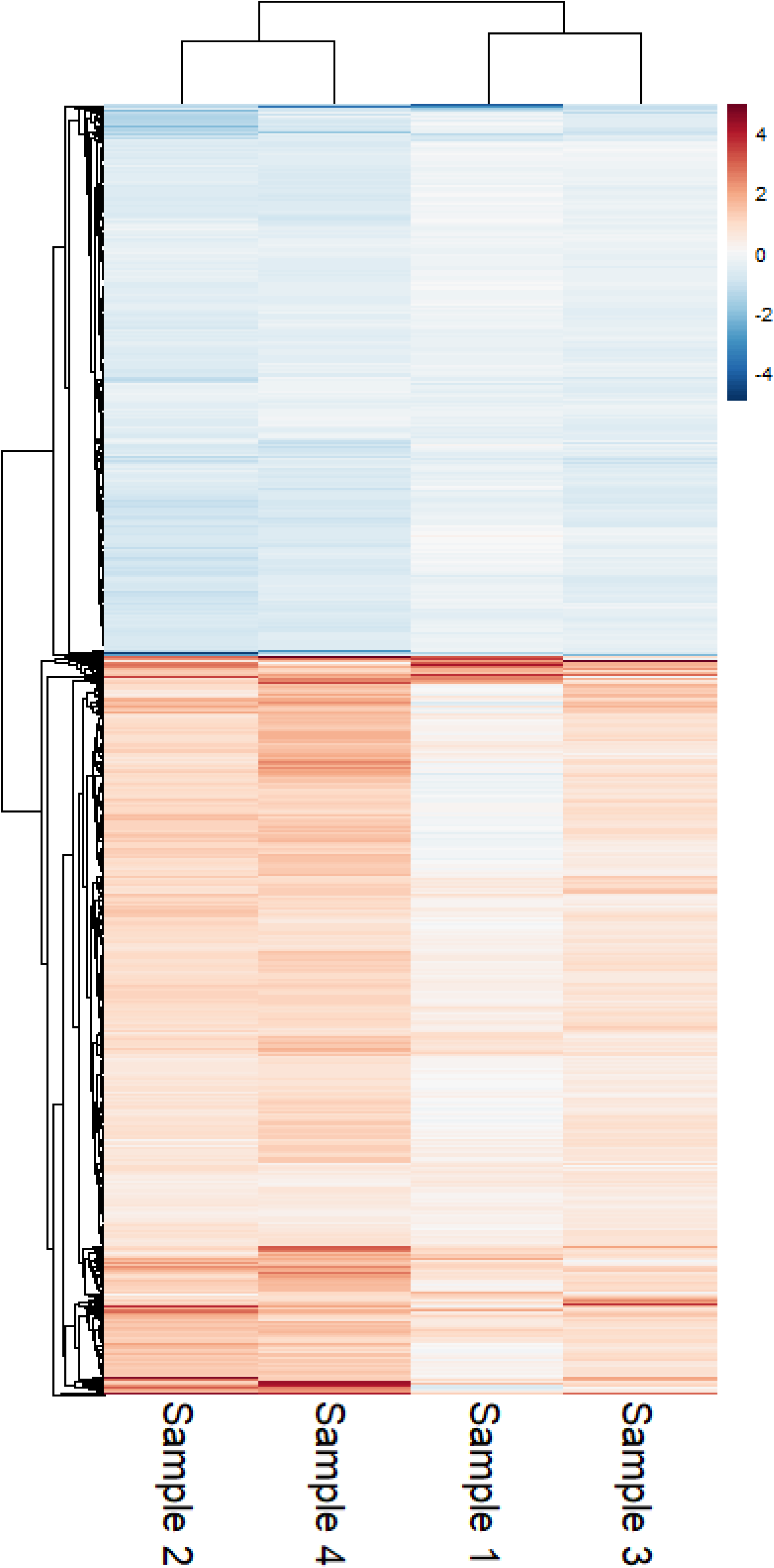
Heatmap of the relative expression of 5894 differentially expressed genes per fish, normalized to the paired control.

For the downregulated genes, 327 enriched GO terms were identified (adjusted *p*-value<0.05) (Dataset 2). The top 15 most significant (adjusted *p-value* 3.90E-36 to 8.33E-13) are shown in Figure 5A, of which 14 were cell cycle-related and one belonging to reproduction. For example, “DNA metabolic process” (177 genes), “Cellular response to DNA damage stimulus” (142 genes) and “Mitotic cell cycle” (140 genes). Furthermore, 15 enriched KEGG pathways were identified (adjusted *p*-value<0.1) (Figure 5B). The most enriched pathway (adjusted *p-value* 7.50E-13) also containing the highest number of genes (61 genes) was “Cell cycle”, a pathway describing mitotic cell cycle progression. To deeply study downregulated reproduction-related genes, we selected the “meiosis II” cascade of the “oocyte meiosis” pathway, which contained 8 downregulated genes: adenylate cyclase 5 (*adcy5*), cysteine three histidine 1 (*cth1*), ribosomal protein S6 kinase a, polypeptide 1 (*rps6ka1*), BUB1 mitotic checkpoint serine/threonine kinase (*bub1*), mitotic arrest deficient 1 like 1 (*mad1l1*), cyclin E1 (*ccne1*), anaphase promoting complex subunit 2 (*anapc2*) and cell division cycle 20 homolog (*cdc20*). Among them, we identified those genes which matched the seed region sequence of miR-210 in their 3’UTR. In total, four genes matched the seed region as shown in Figure 6 —and more in detailed in Supplementary figure S4—, namely *adcy5*, *rps6ka1*, *mad1l1* and *ccne1*, overall indicating that miR-210 was able to downregulate Meiosis phase II.

**Figure 5.**
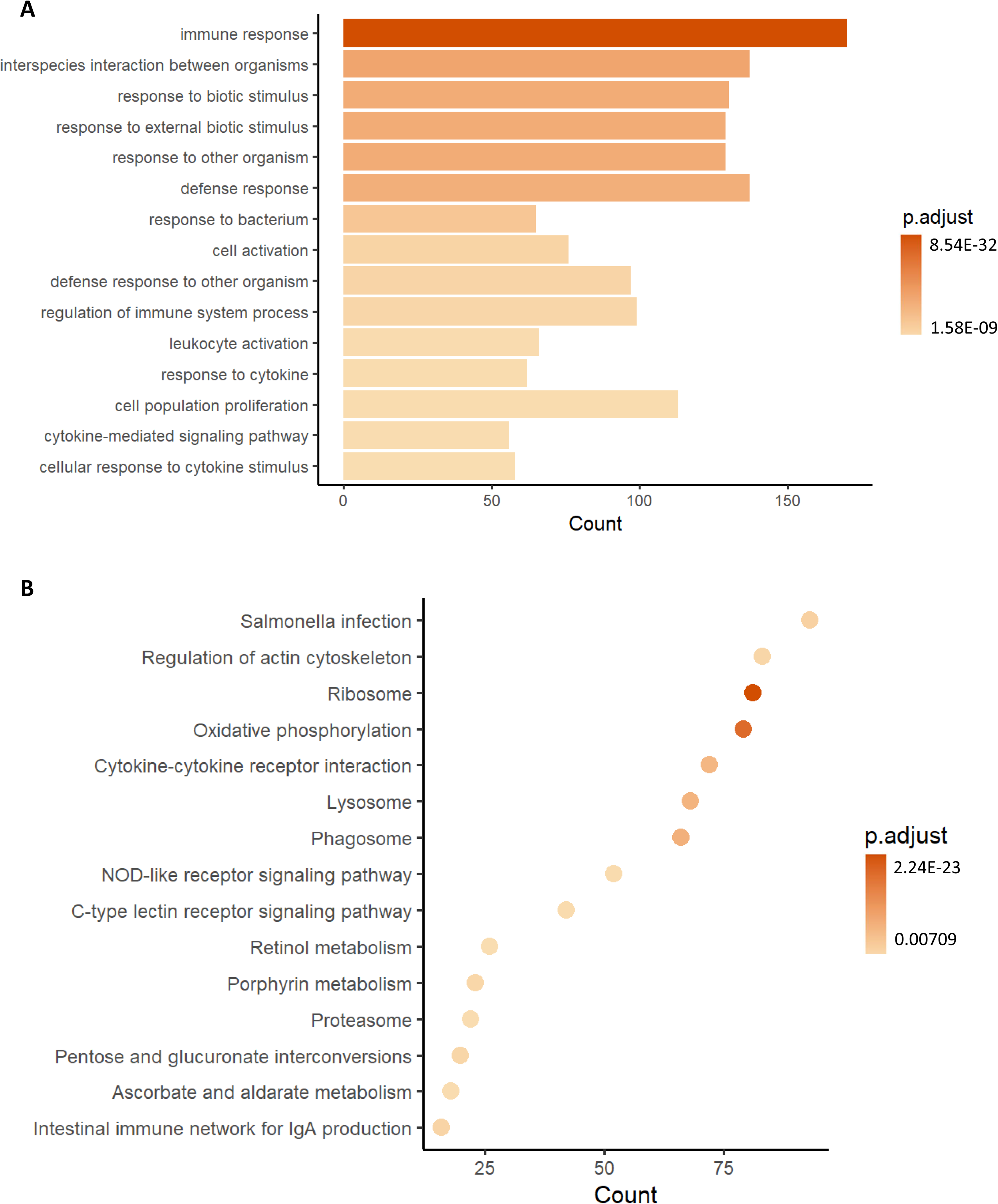
Top 15 enriched GO terms (A) and KEGG pathways (B) of upregulated genes.

**Figure 6.**
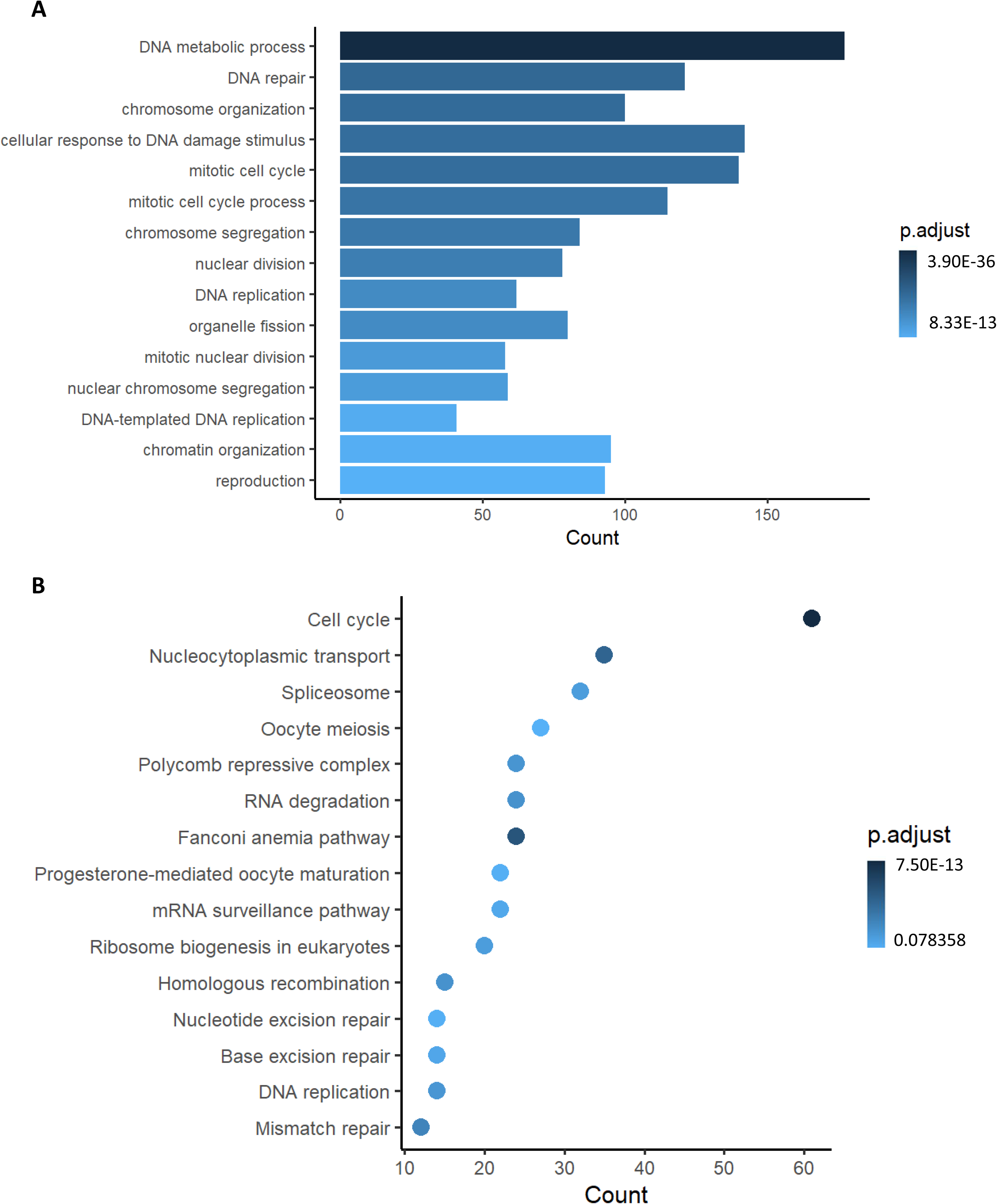
Top 15 enriched GO terms (A) and KEGG pathways (B) of downregulated genes.

**Figure 7.**
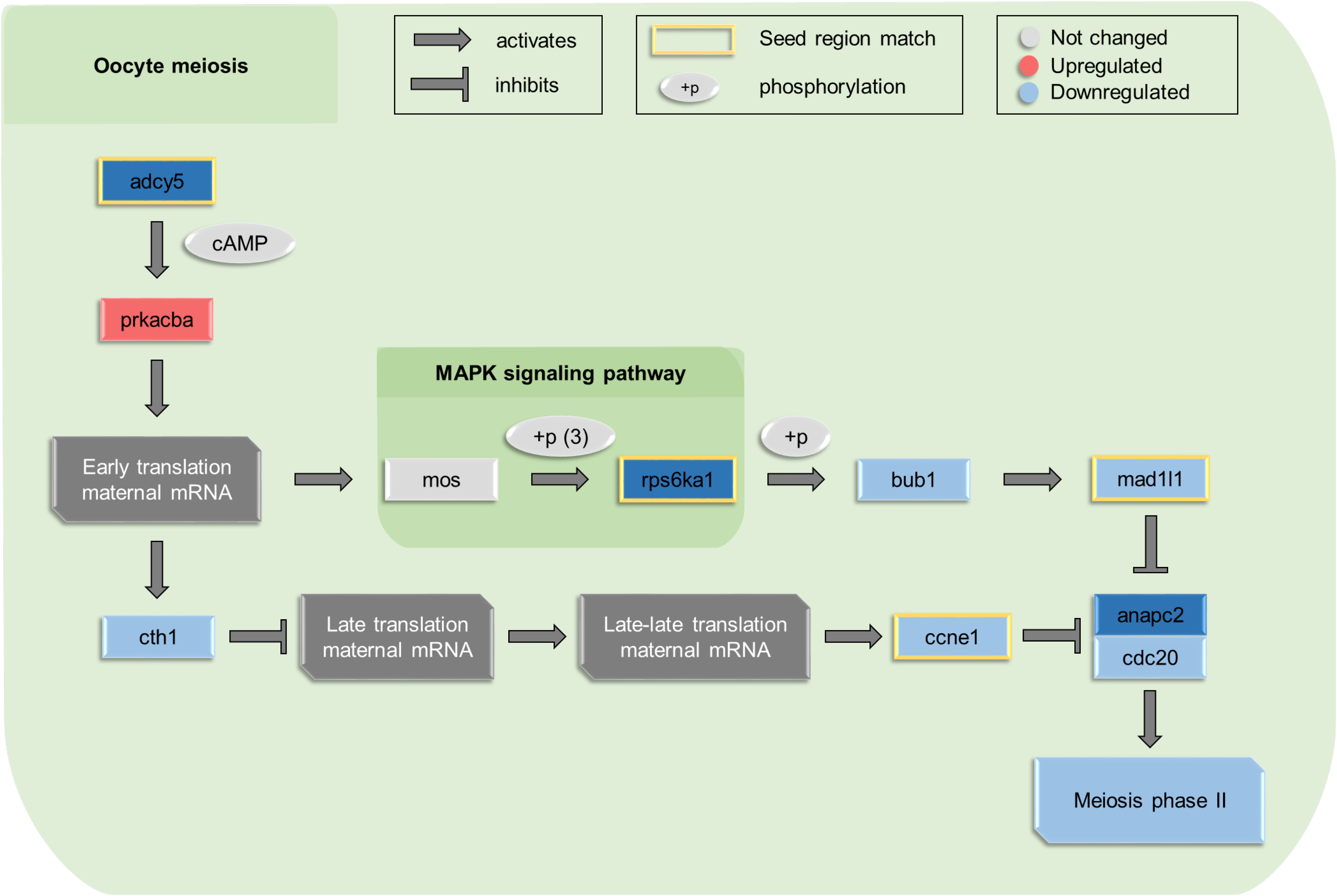
A simplified overview of the oocyte meiosis KEGG pathway. Differentially expressed genes are red (upregulated) or blue (downregulated) in gradients of differential expression. Genes with matching seed regions to dre-miR-210 are marked in yellow.

### In silico cell type identification

DEGs were assigned to cell types based on the marker genes previously described by Liu *et al*. (Liu *et al*., 2022) and Jiang *et al*. (Jiang *et al*., 2021). In total, from our sequencing data, 495 DEGs were assigned to immune cells, 912 to germ cells and 604 to ovarian supporting cells (Dataset 3). Cell types were divided into ten categories: innate immune cell, granulocyte, lymphoid cell, myeloid cell, pre-oocyte, oocyte, somatic cell, vasculature cell, epithelial cell and other (Figure 8A). We were able to classify 49, 86, 75 and 429 DEGs as innate immune cells, granulocyte, lymphoid cell and myeloid cell, respectively, and also 511 and 437 DEGs as pre-oocyte and oocyte markers, respectively. Furthermore, 604, 98 and 114 were identified as somatic cell, vasculature cell and epithelial cell, respectively. Among them, the most upregulated genes were *amh* (follicle and granulosa cell), *postnb* (stromal cell) and *pcolcea* (theca cell).

**Figure 8.**
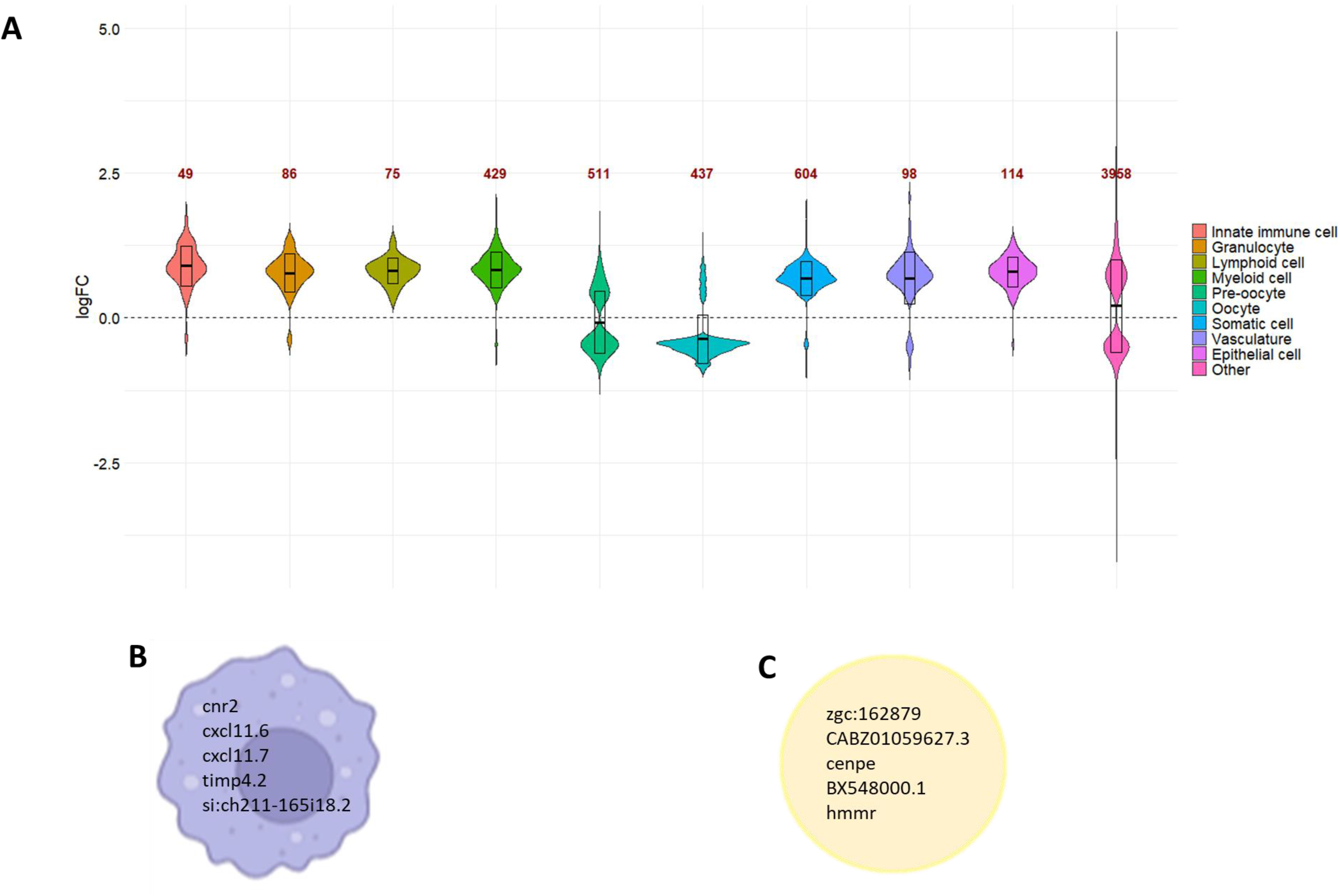
A. Violin plot of differentially expressed genes classified per cell type based on Liu *et al*. and Jiang *et al*. Cell types are divided into immune cells, *i.e.* innate immune cells, granulocytes, lymphoid cells and myeloid cells, ovarian cells, *i.e.* pre-oocyte and mature oocyte, somatic cells, vasculature cells, epithelial cells and undefined (other). The number of DEGs corresponding to each cell type are shown in red text. B. Top 5 upregulated genes in macrophages. C. top 5 downregulated genes in oocytes.

In order to identify reproduction-related gene markers for specifically ovarian cell types, DEGs were further studied in different oocyte maturations (Dataset 3). The top 5 downregulated target-genes in the oocyte were zgc:162879, CABZ01059627.3, centromere protein E (*cenpe*), BX548000.1, hyaluronan-mediated motility receptor (RHAMM) (*hmmr*), (Figure 8B) and the most downregulated were (PGC), *dazl* (PGC), *aspm* (pre-mei GC), *cenpe* (meiosis GC), *ca15b* (post-meiosis GC), *grip2a* (early oocyte).

Similarly, to identify immune-related gene markers in the ovarian cell culture, immune-related markers found in the immune cells such as macrophages, neutrophils, T-cells and natural killer (NK) cells, were identified (Dataset 3). The top5 upregulated target-genes after miR-210 incubation in macrophages were: cannabinoid receptor 2 (*cnr2*), chemokine (C-X-C motif) ligand 11, duplicate 6 (*cxcl11.6*), chemokine (C-X-C motif) ligand 11, duplicate 7 (*cxcl11.7*), TIMP metallopeptidase inhibitor 4, tandem duplicate 2 (*timp4.2*) and si:ch211-165i18.2 (Figure 8C) and the most upregulated were *cnr2* in neutrophil, *lpar5a* in T-cell, and *ncaldb* in NK-cell.

## Discussion

In this study, we showed the downstream effect of dre-miR-210-5p on the ovarian transcriptome in a fish model. Higher expression levels of this miRNA caused an activation in the immune response and inhibited oocyte meiosis together with DNA replication and cell division. Although the existence of the interactions between the reproductive system and the immune system in fish gonads exist, much still remains to be understood. It is of great interest to study the relationships between these two systems, not only to control the vertical transmission of pathogens, but also to understand how fish are able to cope the infections present in the gonads. The gonads are organs with immune privilege, meaning that an immune response can be deferred (Maddocks and Setchell, 1990). In fish, the majority of studies concerning the relationships between these two systems are found in the field of immuno-toxicology because a large number of endocrine disruptors can mimic sex steroids and, in turn, activate the immune system (Weber et al., 2019). The expression of genes related to the immune system during gonadal development has been described in some, although few, fish species. This is the case of gilthead sea bream (*Sparus aurata*), a hermaphroditic teleost which requires a massive infiltration of an immune cell type (acidophilic granulocyte) for the transformation of the testes during sex change (Liarte *et al*., 2007; Chaves-Pozo, García-Ayala and Cabas, 2018), in turbot (*Scophthalmus maximus,* (Ribas *et al*., 2016), in European sea bass *Dicentrarchus labrax*, (Ribas *et al*., 2019) and, in zebrafish (Liew and Orbán, 2014).

Determining direct targets of miRNAs has some bottlenecks. miRNA-mRNA binding depends on free energy GΔ (Ragan, Zuker and Ragan, 2011), the higher the similarity in reverse complement of the sequences, the higher the chance of binding. Techniques such as luciferase assays analyses the binding of miRNA to its 3UTR target by measuring the expression of a fluorescent marker attached to the 3UTR (Tomasello, Cluts and Croce, 2019). This technique, however, does not take the natural structure of the mRNA into account, since only the 3’UTR sequence is synthetized, and thus does not represent an authentic miRNA-mRNA interaction (Hofacker, 2007). Here, we presented a protocol for transfecting miRNA mimic in primary ovary cell culture. A reliable method to study the miRNAs function miRNA transfection protocols have been performed in the established and commercially available ovarian cell line of Chinese hamster ovary cells (Barron *et al*., 2011; Fischer *et al*., 2013; Silveyra, DiAngelo and Floros, 2014) and in the human ovarian cancer cell lines OVCAR3 and SKOV3 (Zhao *et al*., 2021). Alexandri *et al*. (2022) described miRNA mimic transfection on whole mouse ovaries (Alexandri, Van Den Steen and Demeestere, 2022). In Jiang and Ma (2018) (Jiang and Ma, 2018), a primary cell culture of ovarian granulosa cells in rats was established, upon which miRNA mimic transfection was performed. However, until now, no protocol described miRNA mimic transfection on a heterogeneous primary ovarian cell culture in fish. In the current study, we implemented an *in vitro* protocol to study the miRNA downstream effects on the ovarian cell. Our method, based on the miRNA mimic molecules synthetically produced, was successfully validated by: 1) high cell survival 24h after the incubations (average 62% compared to control group); 2) great integrity of the cells studied under fluorescence microscopy; 3) increasing the copy number dre-miR-210-5p in the cells revealed by qPCR analysis; 4) ovarian transcriptome was altered identifying more than 6,000 DEGs; 5) *in silico* analysis revealed that the immune-system was activated while the reproduction system was inhibited. It is worth noting that the transfection reagent stressed the cell culture by damaging and decreasing cell survival. In our protocol, some preliminary experiments were performed to adjust the most convenient transfection reagent volume, finding the proper TR to mimic ratio. Similarly, as previously described, the optimal concentration for RNA mimics is critical for cell cultures success (Jin *et al*., 2015).

In total, we identified ∼6,000 DEGs of which ∼2.500 downregulated and ∼3.400 upregulated genes after 24h of miRNA-210 incubations. Even though miRNAs directly cause mRNA suppression, downstream effects of miRNA overexpression may result in upregulation of transcripts (O’Brien *et al*., 2018). Furthermore, theories of miRNA-mediated upregulation have been described as well (Orang, Safaralizadeh and Kazemzadeh-Bavili, 2014; O’Brien *et al*., 2018). Remarkably, 16 genes have previously been identified as direct targets of miR-210. Among them, four oocyte meiosis-related genes were highlighted as potential target genes based on their differential expression and seed region complement in zebrafish: ribosomal protein S6 kinase a, polypeptide 1 *rps6ka1* (8mer), adenylate cyclase 5 *adcy5* (6mer), Cyclin E1 *ccne1* (8mer) and mitotic arrest deficient 1 like 1 *mad1l1* (7mer-m8).

In our data, we observed an upregulation of genes mainly involved in the immune system, as determined by enriched GO terms and KEGG pathways. Recently, miR-210 played a role in the inflammatory response, where knockout of the miRNA caused limitations in the cytokine storm, decrease cell survival and induced a proinflammatory response in mice (Virga *et al*., 2021). Additionally, activation of the immune system through lipopolysaccharide (LPS) injection caused a significant upregulation of miR-210 in mice macrophages making a suitable diagnostic and prognostic marker for sepsis (Virga *et al*., 2021). On the other hand, most downregulated genes were involved in the cell cycle, in concrete DNA replication, mitosis and meiosis. It was shown that hsa-miR-210 upregulation correlated with suppression of several mitosis-related genes in human, namely Plk1, Cdc25B, Cyclin F, Bub1B and Fam83D (He *et al*., 2013). In our data, we found downregulation of *plk1*, *cdc25b, ccne1* and *bub1bb* after dre-miR-210 upregulation. These genes played key roles the reproduction system, as it was described, for example, that *plk1* was involved in the transition of the G2/M phase in Nile Tilapia gonads (Matsuoka *et al*., 2008). In mice, knockout of *cdc25b* caused infertile females due to meiotic arrest of oocytes (Lincoln *et al*., 2002). Extensive data is found for *ccne1* gene in the reproduction system in many species, as being an oncogenic driver for ovarian cancer (Gorski, Ueland and Kolesar, 2020; Xu *et al*., 2021; Yao *et al*., 2022; Kang *et al*., 2023). In fish, less data is found describing the role of *ccne*. For example, in triploid female rainbow trout (*Oncorhynchus mykiss*) this meiotic gene was targeted by dre-miR-15a-5p during gonadal development (Huang *et al*., 2021) as well as upregulated in the zebrafish liver after toxic insults (Carlson and Van Beneden, 2014).

miR-210 demonstrated distinct actions depending on the specific cell type. With the aim of deciphering downstream gene alterations of the miRNA-210 in the different type of cells of the ovarian cell culture, cellular markers for reproduction-and immune-related cells type were used (Liu *et al*., 2022) (Jiang *et al*., 2021). Related to the immune system, Jia et al. (2019) (Jia *et al*., 2019)described the suppressive role of miR-210 in primitive myelopoiesis by targeting *gata4/5/6* transcription factors, where *gata4/5/6* knockdown resulted in increased miR-210 expression and overexpression of *gata4/5/6* caused a decrease in expression. During satellite cell differentiation in rainbow trout (*Oncorhynchus mykiss*), miR-210 was upregulated (Latimer *et al*., 2017). Moreover, miR-210 inhibits apoptosis in ovarian follicle cells of marine medaka (*Oryzias melastigma)* by targeting apoptosis-genes (Tse *et al*., 2015).

Since miR-210 is hypoxia-induced, in humans, this miRNA is prominently present in the tumor microenvironment, where hypoxia is a common feature and currently used as a marker for cancer diagnosis (Emami Nejad *et al*., 2021). Oncogenic properties of miR-210 were reviewed in Qin *et*. *al.* (Qin, Wei and Li, 2014), where cancer-stimulating processes were described to which this miRNA has been linked, such as angiogenesis, anti-apoptosis and cell proliferation. Several targets of miR-210 functioning as oncogenes were described, of which four genes were downregulated in our data (*casp8ap2*, *e2f3*, *RAD52* and *tp53i11a*) and one upregulated (*hoxa1a*). Furthermore, the diagnostic significance of miR-210 in human cancer was calculated in Feng *et al*. (2019) (Feng *et al*., 2019). By correlating the miR-210 expression with the presence of tumors from various tissues between patients and the control groups, suggesting miR-210 as a biomarker for cancer detection. Specifically, in ovarian cancer, in Li *et al*. (2014) (Li *et al*., 2014), hypoxia-induced miR-210 upregulation promoted epithelial ovarian cancer cell proliferation as well as suppressed in apoptosis, similarly to synthetic miR-210 upregulation as described (Zhao *et al*., 2021).

The presence of (pre-)miR-210 in cancerous conditions in fish was described by using a transgenic melanoma model with the Japanese medaka and natural *Xiphophorus* melanoma model (Mishra, Kneitz and Schartl, 2014). In all tumors, relative expression of miR-210 was upregulated and Hypoxia Inducible Factor 1 Subunit Alpha (HIF1A) was proposed as the target gene (Mishra, Kneitz and Schartl, 2014). miR-210 has been described in several fish species, mainly describing its involvement in the immune system and cell proliferation. In miiuy croaker, miR-210 expression was significantly upregulated in spleen tissue and macrophages after stimulation with poly(I:C), an immune-stimulant used to simulate a viral infection (Sun *et al*., 2018). Here, downregulation of Deubiquitinating enzyme A (DUBA), a negative regulator of IFN signaling, was inversely correlated with miR-210 expression after stimulation (Sun *et al*., 2018). Similarly, miR-210 was expressed in spleen after *Vibrio harveyi* infection and LPS stimulation and was shown to target RIPK2 by inhibiting the inflammatory cytokine production (Su *et al*., 2021). In kidney tissue, miR-210 was involved in the regulation of the rhabdovirus infection (Xu, Chu and Cui, 2018a). Interestingly, miRNA-210 has been described in invertebrate species. This is the case of sea cucumber (*Apostichopus japonicas*) coelomocytes in which miR-210 behaved similarly of that described in the present study because miR-210 increased the expression of the immune system after *Vibrio splendidus* infection while cell proliferation was inhibited (Zhang *et al*., 2020). Furthermore, miR-210 was found in shrimp (*Penaeus vannamei*) infected with white spot syndrome virus (WSSV) (Shekhar *et al*., 2019). These studies unrevealed the importance that miR-210 might have in the immune system throughout evolution.

## Conclusion

miR-210 has been extensively studied in ovarian cancers, but its role in the ovarian cellular environment of zebrafish remains unclear. In our research, we explored dre-miR-210-5p and found that it triggered an immune response while inhibiting oocyte meiosis and cell proliferation. Notably, we successfully conducted a miRNA mimic study in ovarian primary cell cultures for the first time. Our results shed light on the function of miR-210 in fish ovaries, as evidenced by significant changes in the ovarian transcriptome in the zebrafish model. This newfound understanding not only enhances our comprehension of miR-210 in human cancer but also highlights its potential as a marker for various aquaculture-related applications.

## Acknowledgements

This study was supported by the Spanish Ministry of Science and Innovation grant 2PID2020-113781RB-I00 “MicroMet” and by the Consejo Superior de Investigaciones Científicas (CSIC) grant 02030E004 “Interomics” to LR. We thank the lab technician Sílvia Joly for her essential assistance in our team and Gemma Fusté for her assistance in fish facilities. This study was supported by funding from the Spanish government through the ‘Severo Ochoa Centre of Excellence accreditation (CEX2019-000928-S).

## Supplementary info

### Supplementary tables

**Supplementary Table S1.** Primer sequences for qPCR validation

**Supplementary Figure S1.** qPCR miR-210 + MDS

**Supplementary Figure S2.** qPCR validation

**Supplementary Figure S3.** NOD-like receptor

**Supplementary Figure S4.** Full oocyte meiosis

**Supplementary Figure S5.** common genes

### Dataset information

**Dataset 1.** Differentially expressed genes

**Dataset 2.** Enriched GO terms and KEGG pathways

**Dataset 3.** Cell type classification of DEGs

---

**Suppl Fig 1.**
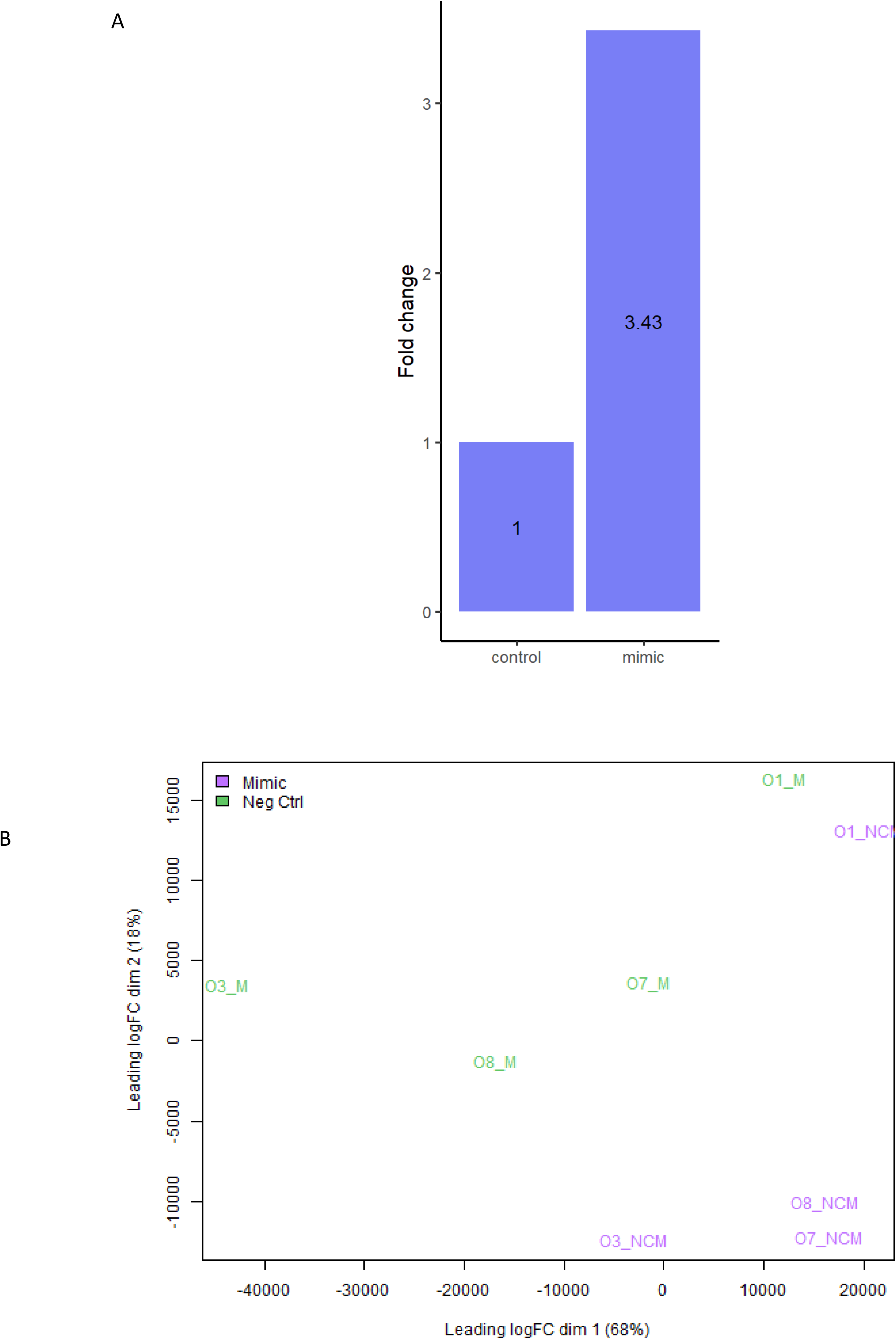
miRNA qPCR + MDS.

**Suppl Fig 2.**
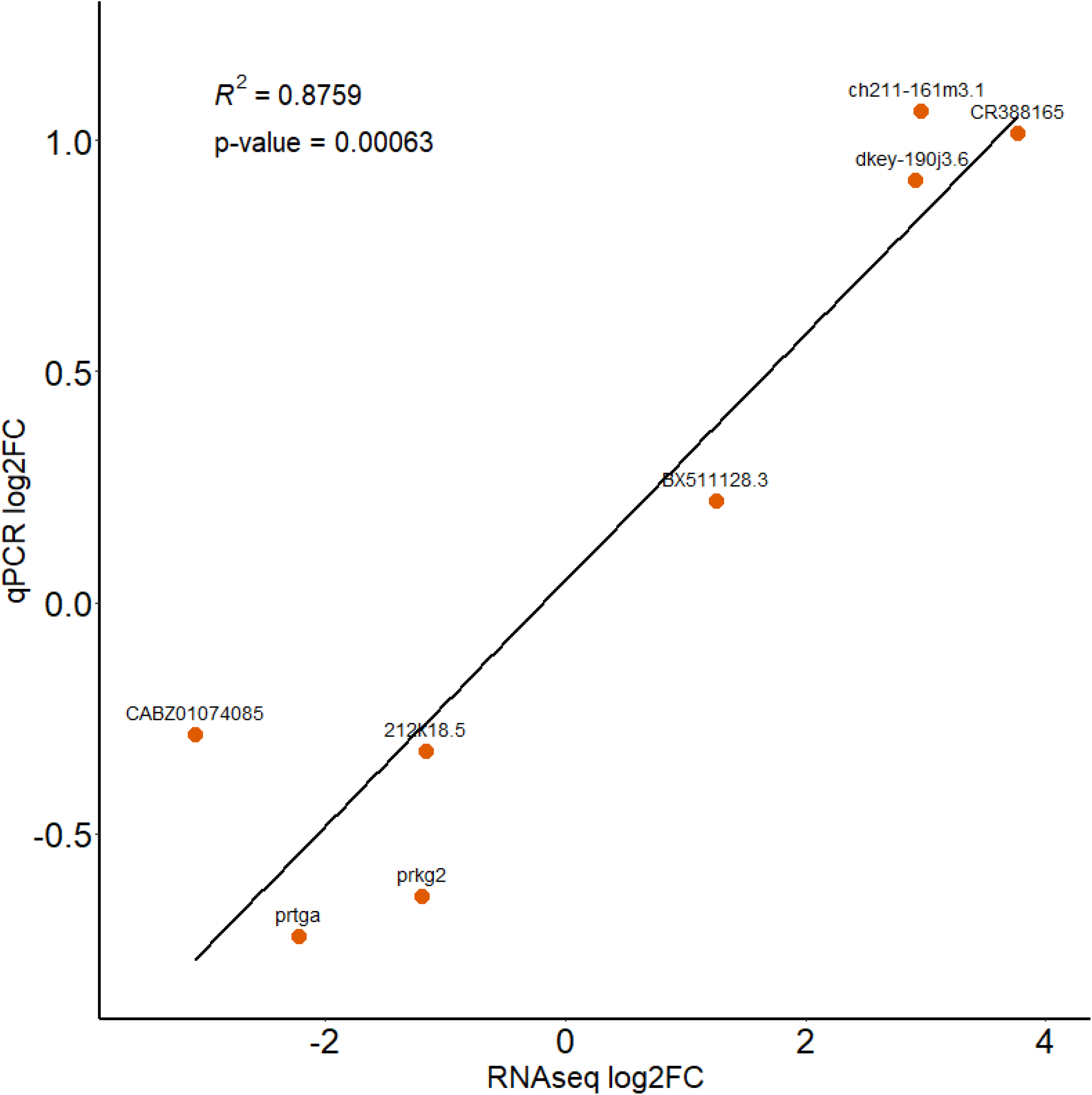
qPCR validation.

**Suppl Fig 3.**
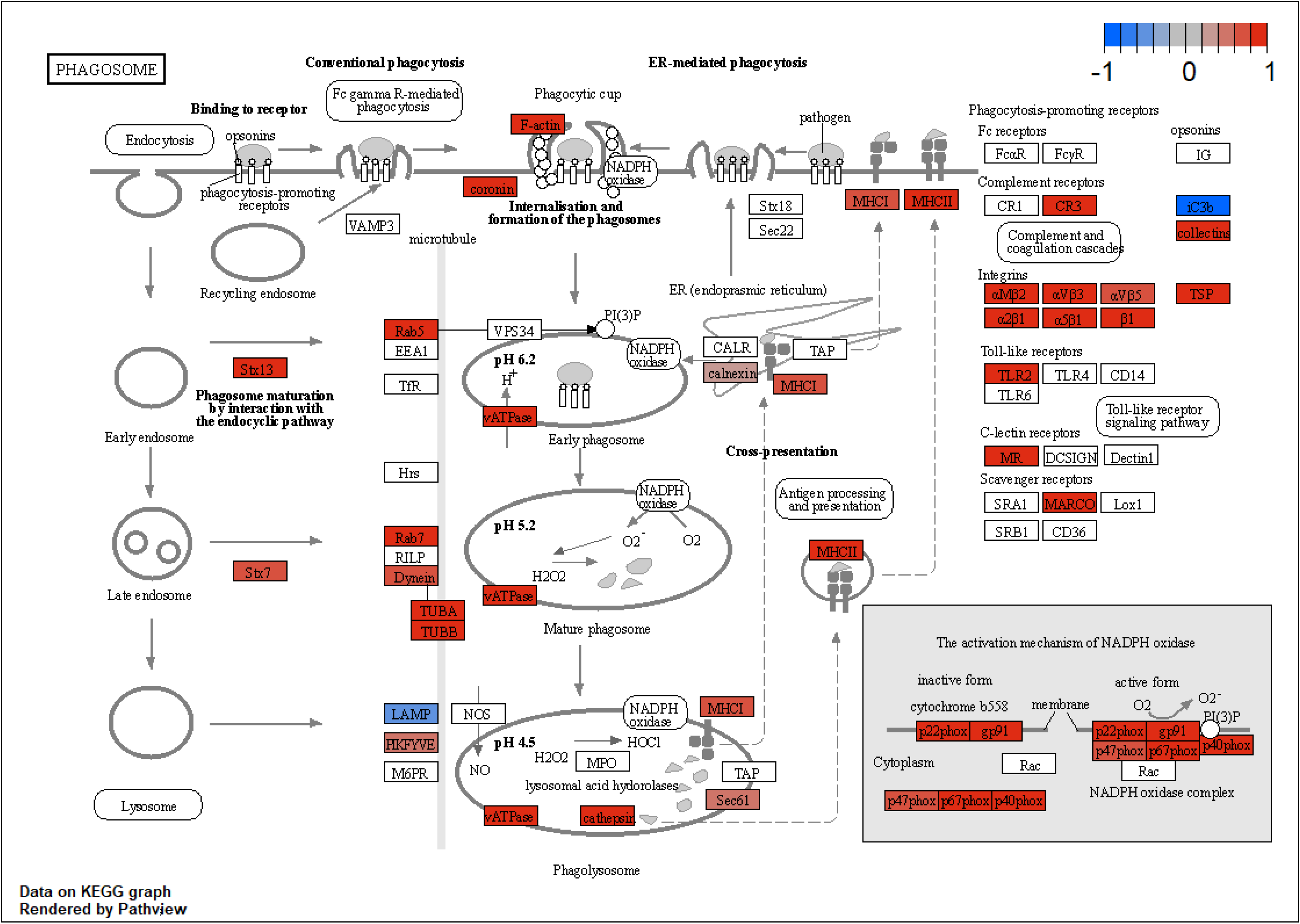
KEGG pathway up.

**Suppl Fig 4.**
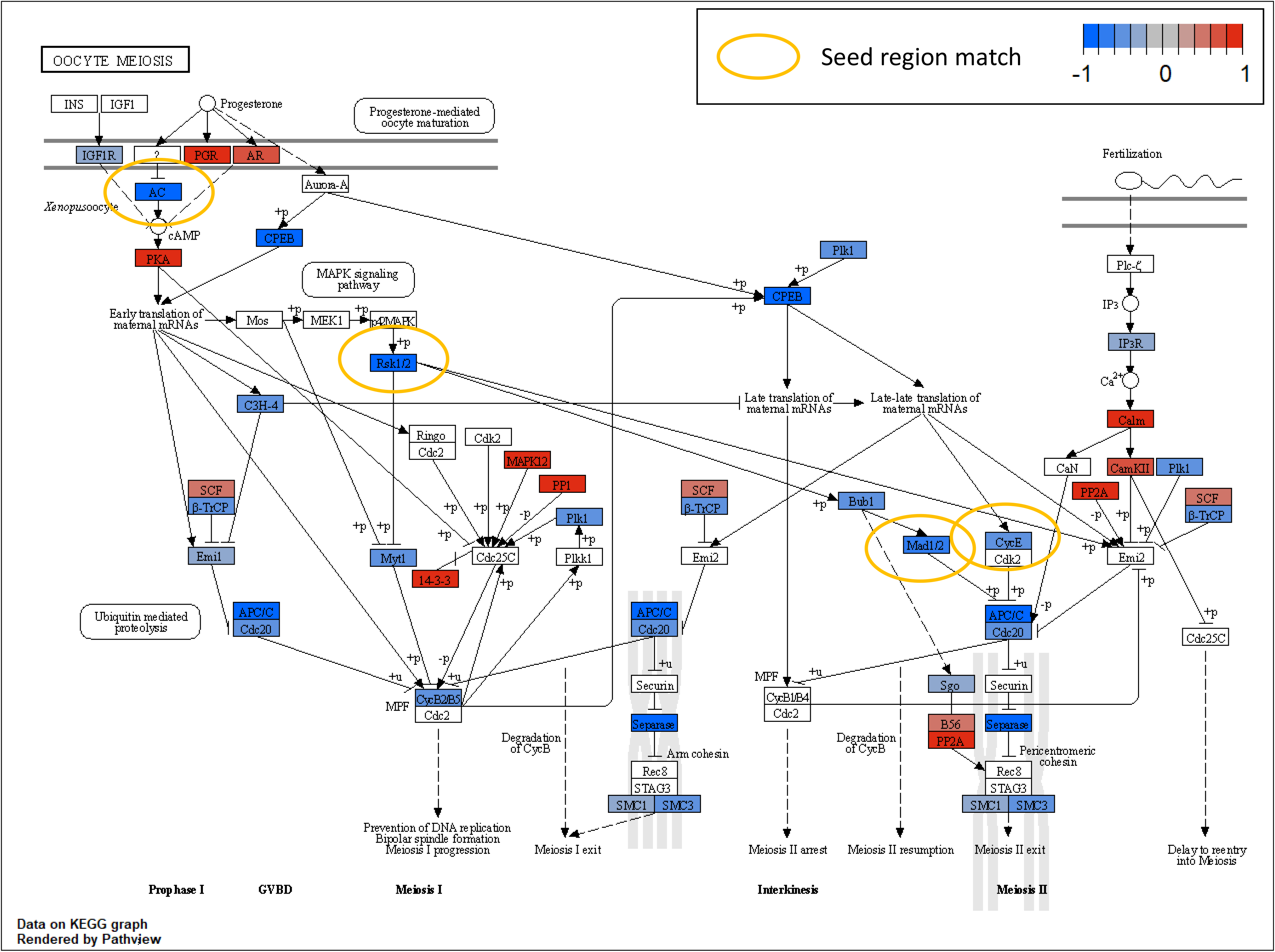
KEGG pathway down.

**Suppl Fig 5.**
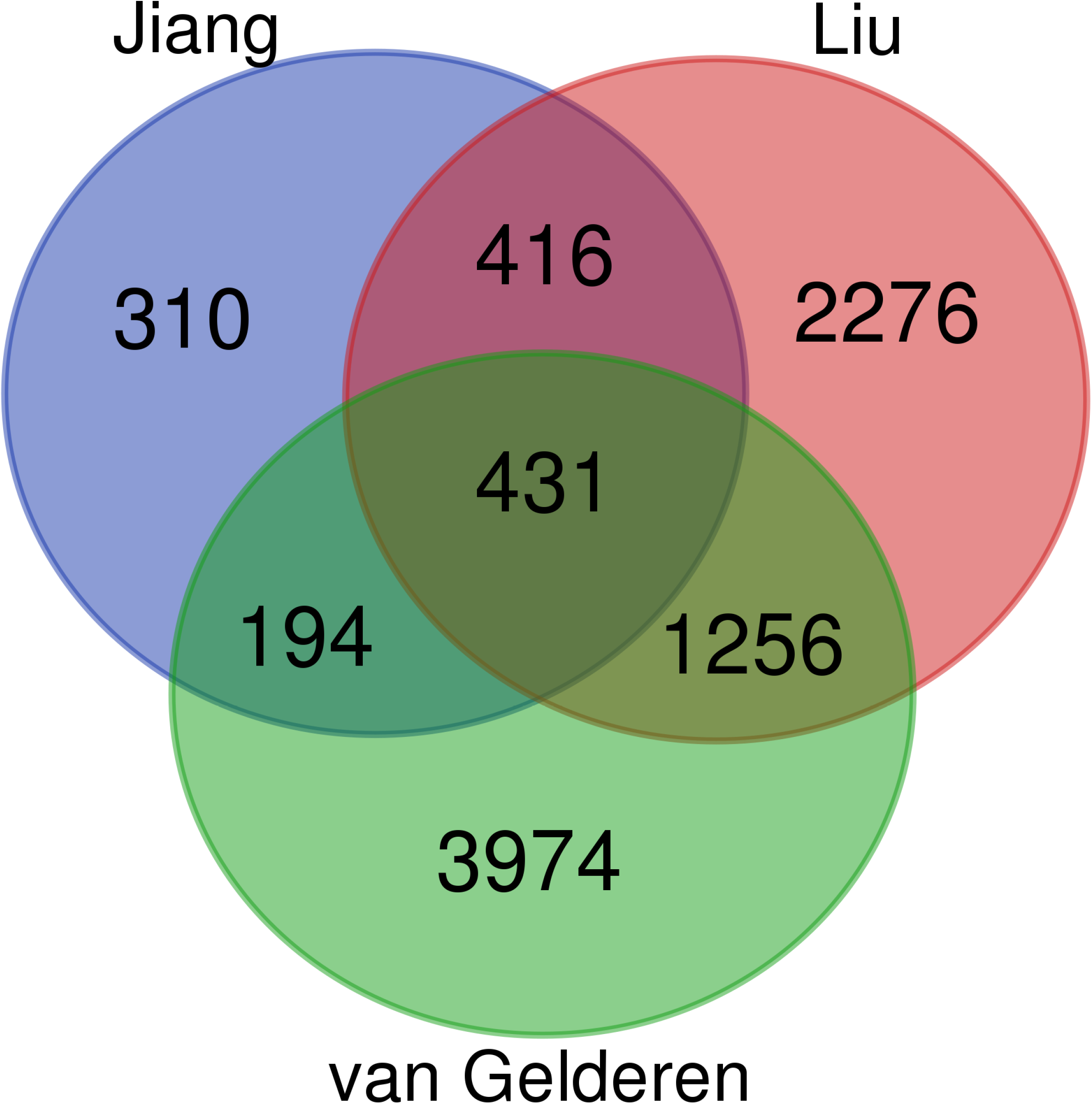
common genes cell types.

